# Rapid evolution and comparative analysis of piRNA clusters in *D. simulans*

**DOI:** 10.64898/2026.01.19.700409

**Authors:** Prakash Narayanan, Satyam Srivastev, Sarah Signor

## Abstract

Eukaryotic genomes are ubiquitously occupied by mobile genetic elements termed transposons, which are silenced via a specialized class of small RNA called piRNA. The small RNA is produced from the transposons themselves when they occupy specialized regions of the genome termed piRNA clusters. The formation of these specialized regions, or their evolution over time, is not well understood. Recent work has suggested that they are extremely variable even within a single species such as *Drosophila melanogaster*. We were interested in taking a comparative approach to piRNA cluster evolution to ask the question - what processes are unique to *D. melanogaster* and which are shared? Shared phenomena are more likely to be fundamental aspects of piRNA formation and evolution compared to those that are more labile. Using five high-quality long-read genome assemblies and five genotype-specific piRNA libraries, we approach this question from a population genetics standpoint. We annotate piRNA clusters, transposons, and structural variants in each of these five genomes. We found extensive variation in piRNA clusters across strains, with smaller piRNA clusters more likely to be limited to a single genotype. By and large, our results are consistent with a model of piRNA cluster evolution in which piRNA clusters are rapidly formed and lost, with a small subset increasing in frequency and length over time. However, we find that the TEs which nucleate the formation of small piRNA clusters are entirely distinct in *D. simulans* compared to *D. melanogaster*, and likely reflect its invasion history rather than any inherent property of the transposon to nucleate clusters. Therefore, while large common clusters can act as ‘traps’ as has been posited for piRNA clusters, there are also numerous small clusters that are born and lost rapidly within a species.

## Introduction

Transposons are mobile genetic elements that introduce a fitness burden to their hosts as they increase in copy number. Transposons can be divided into two major groups - DNA transposons, which replicate themselves in a ‘cut and paste’ manner, and retrotransposons, which use an RNA intermediate to ‘copy and paste’ themselves to new regions of the genome. Transposons are found across the eukaryotic tree of life. Because transposons have deleterious effects, eukaryotes have a dedicated suppression system based on small RNA termed the Piwi-interacting RNA pathway (piRNA) [Brennecke et al., 2007, Houwing et al., 2007]. These small RNAs are 24-30 bp in length and are cognate to the TE that they are intended to suppress [Chen and Aravin, 2021, Houwing et al., 2007]. In many species, including *Drosophila*, piRNA are maternally deposited in the egg, determining the ‘transposon immunity’ of the next generation. piRNAs act as guides for Argonaute class proteins which are able to cleave the transcripts of the TEs, or deposit heterochromatin over their genomic insertions [Darricarrere et al., 2013, Czech et al., 2008, Wang et al., 2015, Le Thomas et al., 2013, Fabry et al., 2021, Batki et al., 2019].

piRNAs are produced from piRNA clusters (piC), regions of the genome that are rich in TE copies that are transcribed into long non-coding RNA [Brennecke et al., 2007]. piCs occupy less than 5% of the genome in species in which they have been well-characterized (primarily flies, mice, and mosquitos) [Brennecke et al., 2007, Houwing et al., 2007, Chirn et al., 2015, Chen and Aravin, 2021]. The structure and function of piCs and their role in TE suppression has diverged among species, but the primary known function of piCs is to suppress transposons. *D. melanogaster* represents the most well-characterized species in terms of understanding the piRNA pathway and piCs. However, it is unknown how generalizable conclusions from *D. melanogaster* have been, in particular because many developmental studies are performed in a single genotype. In addition, available comparisons are largely macro-evolutionary. Comparison on shorter evolutionary timescales is necessary to understand how piCs function and evolve, as macro-evolutionary comparisons are too divergent for inferences based on homologous genes or sequences [Srivastav et al., 2023].

*D. melanogaster* has well-described piCs which we will briefly summarize here. First, we note that *D. melanogaster* has separate piRNA systems for its germline and ovarian somatic cells, and we focus only on germline piCs in this manuscript [Mével-Ninio et al., 2007, Rivera et al., Signor et al., 2023]. piCs are composed of both large (*>*100 kb) and small (*<*10 kb) regions of the genome [Brennecke et al., 2007, Srivastav et al., 2023, Pritam and Signor, 2025]. Small piCs are generally single TE insertions, while large piCs contain many TE insertions and are frequently located at the border of centromeric and telomeric heterochromatin. Both are characterized by an enrichment of young and active TEs, although larger piCs also contain the remnants of older invasions. Initially, it was believed that in *D. melanogaster* large evolutionarily conserved piRNA clusters were the primary drivers of TE suppression, producing upwards of 40% of the total piRNAs [Brennecke et al., 2007]. More recent work has illustrated that not only are these large piCs redundant and potentially dispensable, their involvement in piRNA production is variable [Teixeira et al., Srivastav et al., 2023].

Exactly how piCs evolve to suppress new TEs has been the object of considerable debate over the last decade. In the model originally proposed, the trap model, a TE invades a novel genome and transposes until a copy lands in a piC [Brennecke et al., 2007, Kofler, 2020, 2019, Pritam et al., 2025]. Upon insertion into the piC, the production of piRNA cognate to the transposon is triggered and transposition is suppressed. The trap model has been supported by some evidence – for example, when artificial sequences were inserted into piCs of a host genome, the result was the production of piRNA complementary to the transgene and silencing of reporter constructs [Josse et al., 2007, Muerdter et al., 2012, Srivastav et al., 2023]. Other evidence has contradicted the trap model – for example, when fragments of the *I* -element transposon are inserted into a host genome, it leads to the production of piRNA regardless of whether it inserts into a piC [Olovnikov et al., 2013]. This led to the proposal of the *de novo* model, which posits that piC formation is driven by individual TE insertions and is not sensitive to genomic location [Akulenko et al., 2018, Olovnikov et al., 2013].

Recent work in *D. melanogaster* resulted in the development of a new model that is essentially intermediate between the two aforementioned models [Srivastav et al., 2023]. This ‘birth and death’ model proposes that piCs form frequently in the genome through recent TE insertions [Kofler, 2020, Akulenko et al., 2018, Shpiz et al., 2014, Zhang et al., 2020, Gebert et al., 2021]. Once established, these piCs may increase in size by trapping additional TEs. However, the lifespan of the piC may be short and will be determined by genetic drift and selection. For example, a piC of any size may eventually be eliminated if it lacks insertions from active TEs [Akulenko et al., 2018, Gebert et al., 2021]

Since most piCs have been characterized at macro-evolutionary distances, it is difficult to compare homologous sequences and piC function to understand how they evolve. That is the reason behind the present work - characterizing piCs in *D. simulans*, the sister species of *D. melanogaster*, to understand the dynamics of piC evolution. For example, we will determine if the birth and death model holds in *D. simulans*, and see if we observe the same intraspecific variability in piC expression that has been observed in *D. melanogaster*. This will help us to understand the evolutionary dynamics and function of piCs more broadly, outside the context of a single species.

## Materials and Methods

### Fly stocks

Four *D. simulans* lines *SZ232*, *SZ45*, *SZ244*, and *SZ129* were collected in California from the Zuma Organic Orchard in Los Angeles, CA in February 2012 [Signor et al., 2017a, 2018, 2017b]. *LNP-15-062* was collected in Zambia at Luwangwa National Park by D. Matute and provided to us by J. Saltz (J. Saltz pers. comm., [Matute and Ayroles, 2014, Schrider et al., 2018]. All stocks were maintained on standard Bloomington cornmeal medium at 22 °C under a 12-hr day/night cycle.

### Small RNA library and sequencing

Ovaries were dissected from each of the five genotypes of *D. simulans*. Two replicates were collected two years apart for each of the five genotypes, with 25 adult female ovaries for each replicate. The flies were yeast-fed for two to three days prior to ovary collection. The collected ovaries were stored in TRIzol (Invitrogen). The RNA extraction was conducted using a standard TRIzol protocol. The samples were sequenced by BGI using DNBseq (BGI, Wuhan) and deposited under PRJNA913883 in NCBI SRA [Signor et al., 2023].

### Small RNA processing

For an overview of our approach to piRNA mapping and cluster calling, please see fig. 1. The small RNA reads were mapped to a library of non-piRNA small RNA species (rRNA, tRNA, siRNA, snoRNA, etc.) to remove non-target small RNA with bowtie (v. 1.2.3) and the following parameters (-v 1 -p 1 -S -a -m 50 –best –strata, [Langmead et al., 2009]. Unmapped piRNA were retained and filtered for reads between 23-29 nt with fastp [Chen et al., 2018].

**Figure 1:**
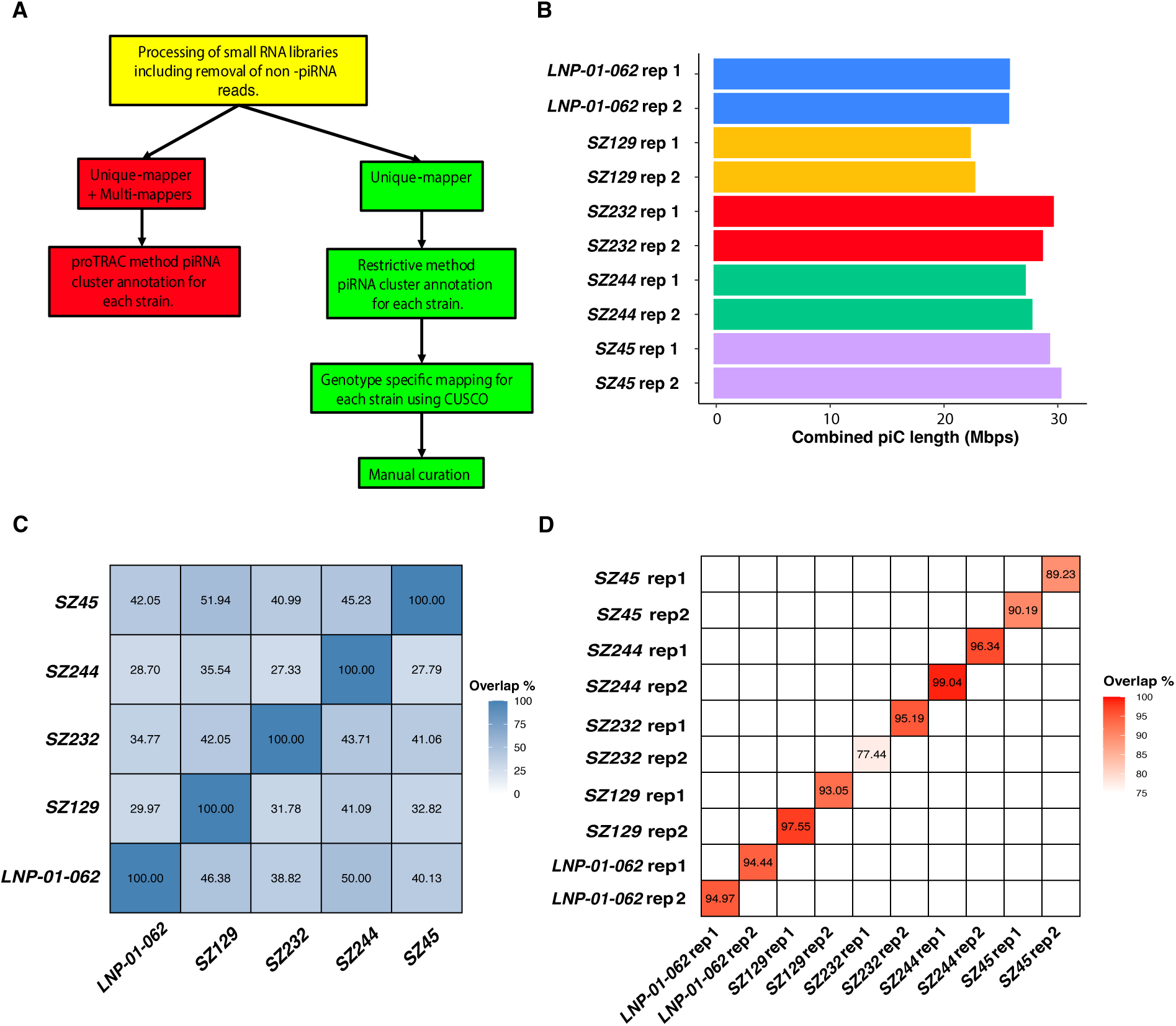
Approach to piC calling and validation of piC position (A) piC calling pipeline (B) Combined piC size predicted for each genome assembly (C) Overlap between strains of predicted piCs identified from each genome assembly. For example, if *SZ45* is used to generate piC positions, 42.5% of those piCs are shared with *LNP-01-062* when the flanks are mapped to the genome of *LNP-01-062*. These comparisons were generated with the restrictive piC calling method. (D) Replicate piRNA libraries were generated two years apart, and the overlap between piC positions of the two replicates is shown here. For example, *LNP-01-062* has a 95% overlap between the piCs called in each of its replicates.

### Genotype-specific genomes

Each piRNA library was matched with a genotype-specific genome assembly. For more information on the methods used to generate the genome assemblies, please see [Signor et al., 2023], and the outline of our methods in fig. 1.

### piC annotation

piC annotation was performed using two approaches, which we will refer to here as the ‘restrictive’ [Srivastav et al., 2023] and ‘proTRAC’ [Rosenkranz and Zischler, 2012] methods. For the restrictive method of annotating piRNA clusters, the piRNA library from each strain was collapsed to unique piRNA sequences using TBr2 collapse.pl (proTRAC) and mapped to the strain’s genome using bowtie ( -n 1 -l 12 -a –best –strata –quiet -y -S) [Rosenkranz and Zischler, 2012, Langmead et al., 2009]. Coverage was then calculated in 500 bp windows with bedtools and converted to RPM [Quinlan and Hall, 2010]. The windows with an RPM value higher than eight were filtered and adjacent windows were merged if the gap between them was less than 20 KB. The clusters were then further filtered for an RPM above 25. Liftoff was used to remap protein-coding genes from the reference genome assembly (Dsim-all-r2.02) to the genome of each of the *D. simulans* genotypes [Shumate and Salzberg, 2021]. Clusters that overlap protein coding genes were split [Neph et al., 2012]. For the proTRAC method, proTRAC was run on this dataset using the following parameters: pdens .07 −1Tor10A .33 −1Tand10A .25 -distr 1-90 -clsize 1000 -clstrand .25 [Rosenkranz and Zischler, 2012]. The resulting clusters were filtered out if the multi-mapped piRNA coverage was 25 or less. Clusters within 20 kb of one another were again merged and split based on the presence of protein coding genes. The restrictive and proTRAC approaches follow that of [Srivastav et al., 2023] but are generally more conservative with RPM cutoffs and cluster size.

### Identifying shared piC regions

The genotype-specific genome assemblies do not have identical coordinates, nor do they have identical piCs. Therefore we had to have a method for identifying shared piCs between different genomes. We chose to identify shared piCs using the flank design portion of the CUSCO pipeline (a method for identifying and evaluating piCs in different genotypes) [Wierzbicki et al., 2023, Wierzbicki and Kofler, 2023]. The flank design portion of the CUSCO pipeline is an approach for identifying homologous piRNA clusters that relies on locating flanking sequences from a reference genome and remapping them to the target genome. We excluded clusters located on unplaced scaffolds (i.e. not on 2L, 2R, etc). Flank design was performed reciprocally for all five genomes – for example the piRNA clusters found in *LNP-01-062* were used to generate flanking sequences and these flanking sequences were mapped to *SZ244*, *SZ232*, *SZ129*, and *SZ45*. Following this, the piRNA clusters that were previously annotated *de novo* with piRNA in each of these genotypes were compared to those uncovered with flank design using bedtools [Quinlan and Hall, 2010]. We required an overlap of at least 20% between piRNA clusters to consider them the same cluster. This percentage was chosen through manual observation of cluster overlap and TE synteny within the cluster.

### Manual curation of piCs

After genotype-specific mapping using the flank design step from CUSCO, the piCs for each strain were individually examined for missing homologous clusters, i.e. a cluster that was mistakenly not identified using the flank design approach. This most commonly occurred because one of the two piC flanks did not map. If one flank mapped and at least three TEs had conserved synteny this was considered a shared cluster.

### Structural variation detection and filtering

Raw long reads for five *D. simulans* genotypes were mapped to the *w^501^* genome (GCF 016746395.2) with minimap2 -N3 and the resulting sam file was converted to bam and sorted [Li, 2018, Li and Durbin, 2009, Signor et al., 2023]. Three structural variant callers - sniffles, cuteSV, and svim - were used for detecting structural variants [Jiang et al., 2020, Sedlazeck et al., 2018, Heller and Vingron, 2020]. All three structural variant callers were run with default parameters along with a minimum MAPQ score of 50, minimum read support of five and minimum structural variant size requirement of 30 nt. The results from all three callers were merged using SURVIVOR (merge vcf 30 2 1 1 0 30) separately for each strain, and any structural variant supported by two callers was retained [Jeffares et al., 2017]. Each *D. simulans* strain was then genotyped for all of the structural variants from the merged files using cuteSV ( –max cluster bias INS 100 –diff ratio merging INS 0.3 –max cluster bias DEL 200 –diff ratio merging DEL 0.5 –min read len 500 –min support 5 –min size 30 –genotype). The genotyped calls for each strain were then filtered to remove inversions, duplications, complex, and imprecise variant calls. Structural variants within 30 nt of each other were collapsed to a single call using bedtools [Quinlan and Hall, 2010]. All of the piRNA clusters for each strain called with the restrictive method were remapped to *w^501^* using the flank design portion of the CUSCO pipeline [Wierzbicki and Kofler, 2023]. Bedtools intersect was used to find the simple structural variant calls overlapping the piCs of each strain.

### Estimating TE enrichment and age in common and unique piCs

The initial step for TE enrichment analysis is annotation of the genotype-specific genome assemblies by Repeatmasker (-pa 16 -a -gff -no is -nolow) [Smit et al., 2013-2015]. Only annotations longer than 500 bp and less than 20% diverged from reference library were retained. We also filtered out TE insertions that were annotated as associated with ribosomal genes (R elements) and satellite elements [Eickbush and Eickbush, 1995]. We extracted the TE annotations that fell within unique and common piC regions for each of the five genotypes of *D. simulans* [Quinlan and Hall, 2010]. We generated ‘expected’ TE frequencies for each type of cluster using 1000 random shuffles and performed a binomial test to compare observed and expected frequency (https://github.com/4ureliek/TEanalysis). We used the output of this analysis to calculate the fold change of TE enrichment for each TE. TE divergence from the reference was used as a proxy for age, which was extracted from the Repeatmasker files.

## Results

### piC calls are highly reproducible

The small RNA libraries were analyzed using the pipeline outlined in (fig. 1A). Briefly, piCs were defined by expression of 23-29 nt small RNAs that overlap TE insertions. In calling these piCs we used two methods, restrictive and proTRAC [Rosenkranz and Zischler, 2012] (Supplemental file 1,2). A key difference between these approaches is that proTRAC incorporates information from multi-mapping reads while the restrictive method does not. We discontinued the proTRAC method because both methods, on average, had 81% overlap for all high-confidence piC calls (piRNA mapping greater than 25 RPM) (Supplemental file 3). The pipeline is described in more detail in the methods section. The proportion of the genome occupied by piCs averaged from 13.75 Mb to 16.35 Mb (fig. 1B). Whether an area is considered a piC is sensitive to the choice of threshold, however we used more stringent parameters than a recent analysis of *D. melanogaster* where 4.8MB to 6.3MB of the genome was considered a piC [Srivastav et al., 2023]. This suggests that a greater proportion of the genome is functioning as a piC compared to *D. melanogaster*. Furthermore, the reproducibility of piC mapping was determined by calling piCs from two replicates collected more than two years apart for each of the five genotypes. After following the same pipeline as described above, any cluster that overlapped by 20% or more was considered the same cluster. There was an average overlap between piC replicates of 92.74%, indicating that the restrictive method yielded highly reproducible piC annotation across all genotypes (fig. 1D, supplemental file 5). This also indicates that within a genotype piCs are a stable attribute of the genome.

### Genotype specific mapping reveals a large number of unique piCs

It was readily apparent as we analyzed piCs in each genotype that many were present in a single strain. We wanted to quantify the frequency of piCs across genotypes using each genotype as a ‘source strain’ for the master list of piCs. Approximately 30 % of the clusters from each strain could not be mapped in the other four strains. We required an overlap of 20% for a piC to be considered the same, which is more stringent than previous approaches but was necessary given the frequency with which we observed minimally overlapping piCs without conserved synteny in TE content. For a piC to be considered ‘homologous’ it must have a common origin and thus share a minimum of 3 syntenic TEs across genotypes. Inter-strain comparisons revealed that on average only 38.61% of the total piCs annotated independently were consistent between genotypes (fig. 1C). Overall, these results indicate considerable variation in the piC landscape across genotypes. We found a total of 812 clusters that were unique and only present in 1 of the 5 genotypes. From our reciprocal mapping of piCs, we estimate that on average 108.6, 49, 27.8 and 87.8 piCs are common to two, three, four, and five genotypes respectively. There is a general downward trend in count as the number of genotypes sharing a piC increases, except for when the piC is shared by all genotypes. This suggests that if a piC increases in frequency it typically goes to fixation rather than remaining polymorphic.

We examined the relationship between the number of genotypes that share piCs and the median length of a piC (fig. 2B, supplemental file 4, Figure. 1). If a piC is unique to a single strain, its median length was 4.5 kb. If all of the sampled genotypes shared a piC, the median length was 43.6 kb (fig. 2B). In general, there is a significant upward trend between the size of a piC and the number of genotypes sharing the piC (R^2^ = 0.784, *p*-value = 0.046). A hypothesis would be that larger piCs are more evolutionarily stable. However, the evolutionary stability of the piC does not necessarily mean that activity level is conserved, as will be discussed below [Srivastav et al., 2023].

**Figure 2:**
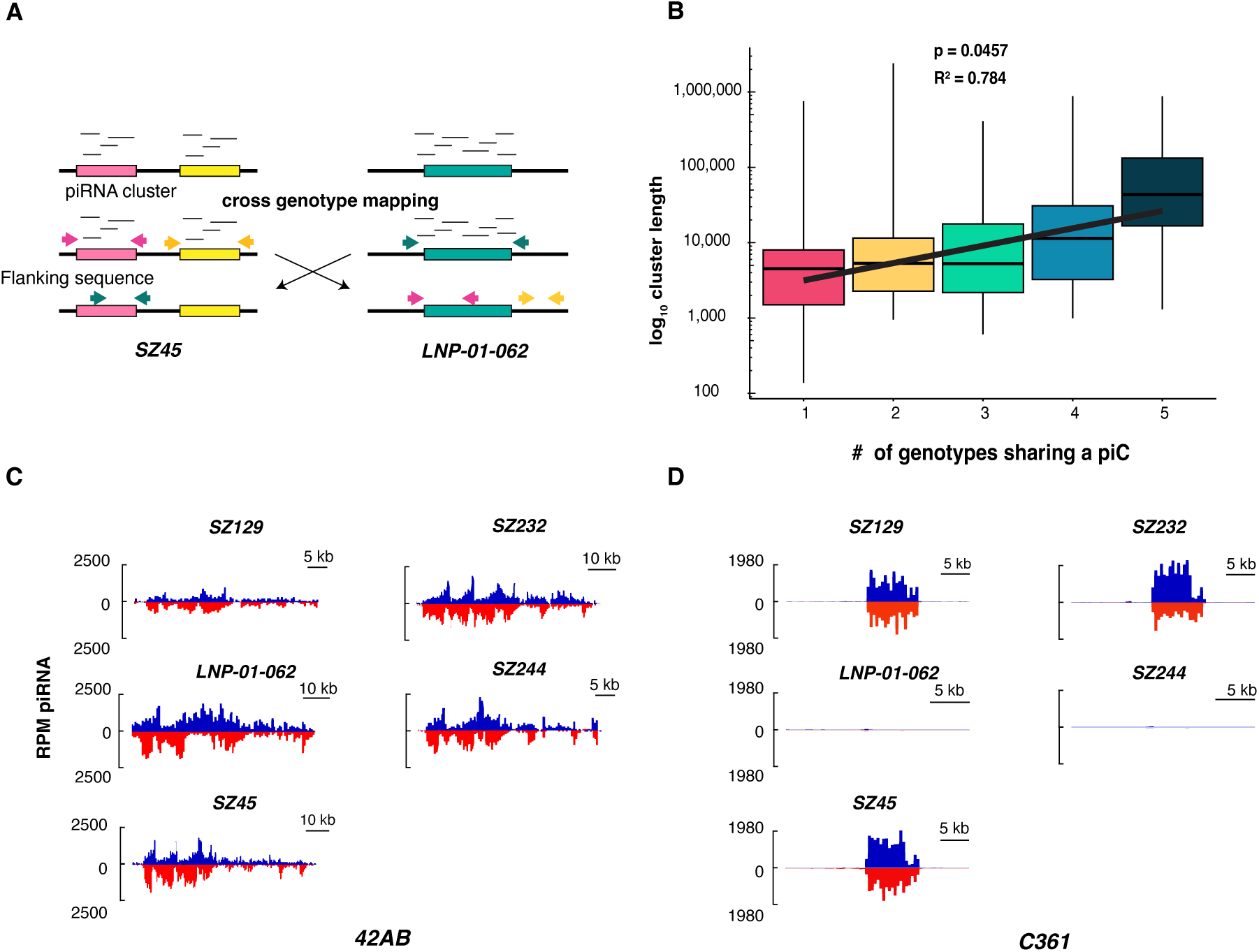
Summary of approach and length distribution of piCs. (A) Each genotype was used to call piCs. The flanking sequences were identified for those piCs, and they were mapped to each of the other four genotypes. This was done using every genome as a reference. (B) Average piC length distribution separated by the number of genotypes shared by a piC. (C) piRNA expression profiles of 42AB for the 5 genotypes. Expression values are in reads per million (RPM). (D) piRNA expression profiles of C361 for the 5 genotypes. Expression values are in reads per million (RPM).

### Variability in piRNA expression of piCs

Recently, Srivastav et al. [2023] demonstrated that one of the major piCs in *D. melanogaster*, *42AB*, was not expressed in every sampled strain. When *42AB* was originally described, 20.8% of total piRNA production was attributed to this locus [Brennecke et al., 2007]. In *D. simulans*, *42AB* is expressed in every strain, however it produces less than half a percent of the total unique piRNA (fig. 2C). This does not place it among the top piRNA producing regions. When the top 15 clusters for each genotype are compared, 8 of them are shared across genotypes, suggesting considerable variation in the contribution each cluster has to the total piRNA pool across genotypes. In each genotype, approximately 62.54% of the total unique piRNAs come from the top 15 unique piCs. The cluster that produces the most unique piRNAs on average (11%) among all genotypes was located on a contig (supplemental file 6).

### Structural variation in piCs

Even if a piC is present in all strains, it can differ considerably in its TE content due to novel insertions and deletions. We wanted to understand the insertion/deletion dynamics of TEs within piCs by examining the patterns of inter-strain structural variation (SV). We detected SVs genome-wide for each strain relative to the *w^501^* reference strain. We mapped long reads from each strain to the *w^501^* reference strain and called SVs using three independent SV callers (see Methods). The SVs supported by at least two callers were merged and genotyped in each strain. The retained insertion counts ranged from 3516 to 4918 and the deletion counts ranged from 3296 to 4551 relative to the reference genome (Supplemental file 4, Figure. 2). On average, 13.6% of insertions and 11.8% of deletions overlapped with the piC calls from their respective genotypes (fig. 3A).

**Figure 3:**
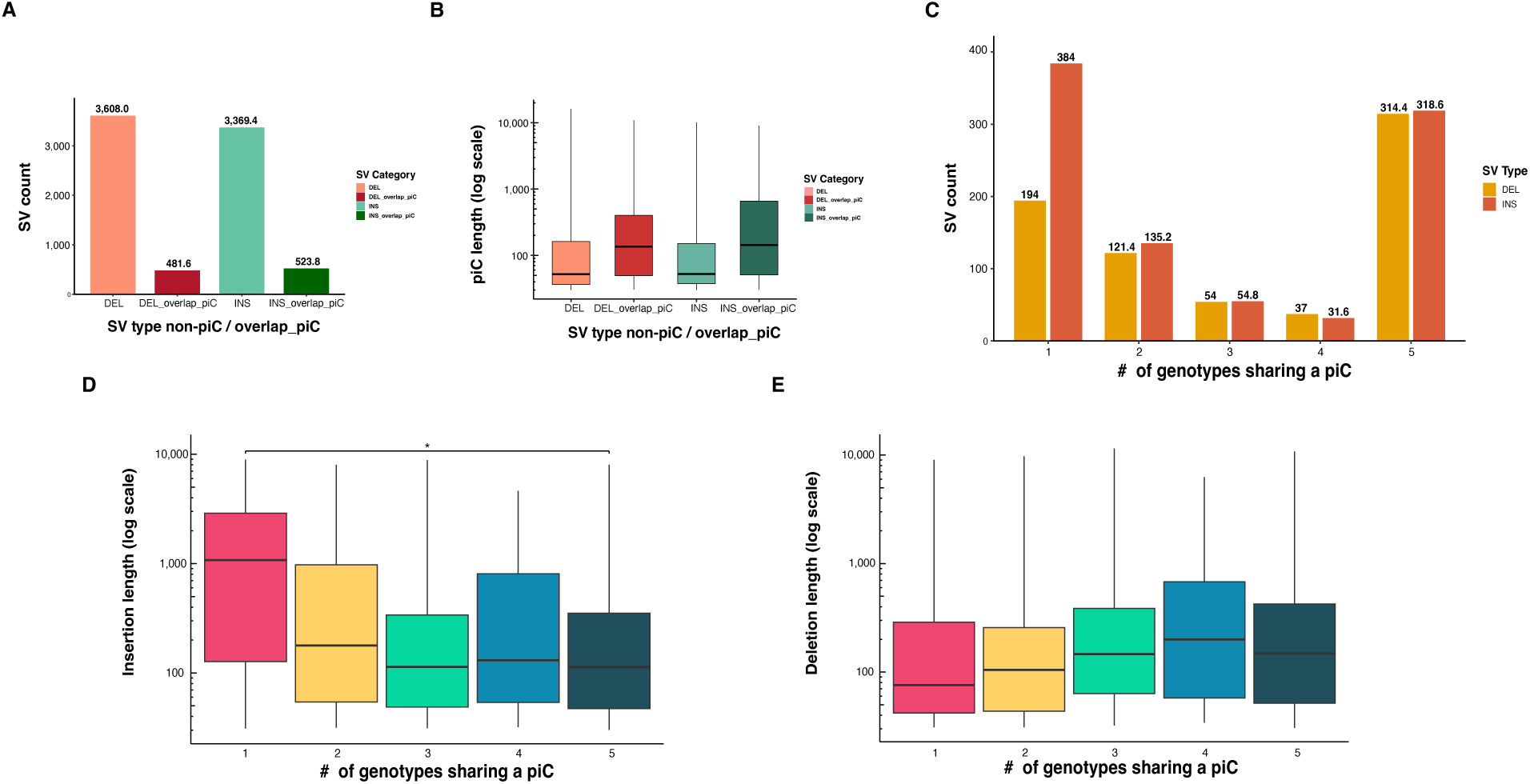
SV distribution in piCs. (A) Average SV counts distribution of insertions and deletions between piC overlap and non-overlap. (B) Average SV length distribution of insertions and deletions overlapping and non-overlapping with piCs, quantified for each strain. (C) Average count of insertions and deletions overlapping piCs grouped by the number of genotypes they are shared by, quantified after remapping using CUSCO to genome assembly of each of the five genotypes acting as reference genome. (D) Average length distribution of insertions overlapping with piCs grouped by the number of genotypes they are shared by, quantified after remapping using CUSCO to genome assembly of each of the five genotypes acting as reference genome. (E) Average length distribution of deletions overlapping with piCs grouped by the number of genotypes they are shared by, quantified after remapping using CUSCO to genome assembly of each of the five genotypes acting as reference genome.

We examined the size distribution of SV types when present within piCs and in non-piC regions of the genome for each strain (Supplemental file 4, Figure. 3). Within piCs the average median length of a deletion was 136 nt, while in non-piC regions it was 52 nt. The largest deletion from a non-piC regions was 16 KB, while in piC regions it was 11 KB. The average median length of insertions also increased in piCs, from 52 nt in non-piC regions to 143 nt in piCs (fig. 3C, Kruskal Wallis p *<*.05). The largest insertions in piC and non-piC regions was similar, at approximately 10 KB. Next, we looked at the frequency of indels in common versus unique piCs. Unique piCs in all genotypes had a larger proportion of insertions (66.4%) compared to all SV types in common piCs (50.3 %)(fig. 3C, supplemental file 4, Figure. 4).

**Figure 4:**
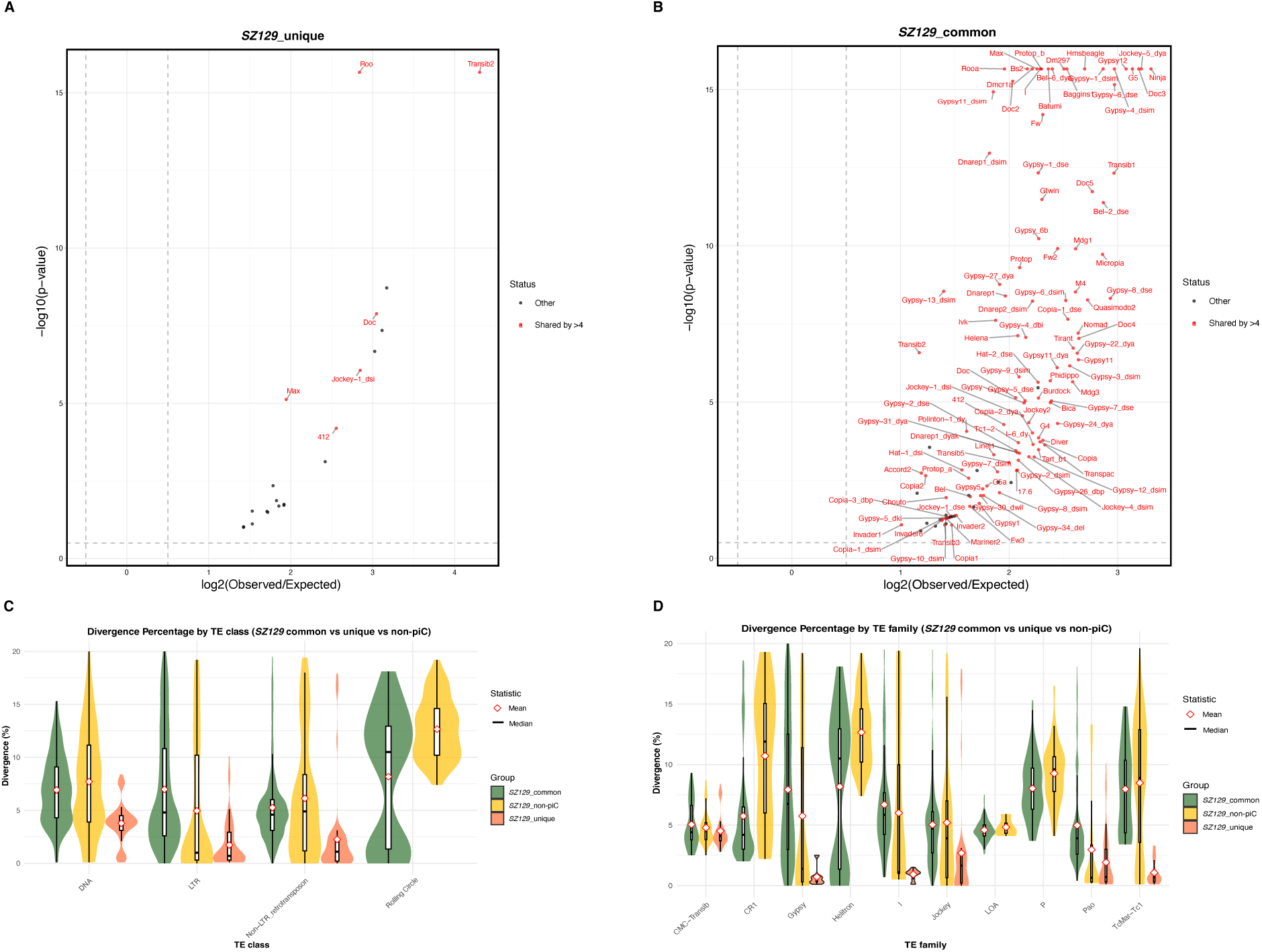
(A) Enrichment of TEs in unique and common piCs of *SZ129*. (B) Enrichment analyses of TE class in unique and common piCs of *SZ129* strain using random shuffling. P -values on y-axis are from binomial test conducted to compare observed overlap counts to expected average overlaps of de novo TE insertions to piCs for each family. (C) Divergence percentage distribution of TE classes in common piC, non-piC, and unique piCs of *SZ129* strain. Y-axis is the divergence percentage, and the range is 0-20 %. (D) Divergence percentage distribution of top 10 TE families in common piC, non-piC, and unique piCs of *SZ129* strain. Y-axis is the divergence percentage, and the range is 0-20 %.

Next, we compared the length of indels in unique and common piCs (Supplemental file 4, Figure. 5, 6). The median length of insertions for unique clusters was 1.78 KB, while common clusters had median insertion lengths of only 113 nt (fig. 3D, Kruskal Wallis p *<*.05). In contrast, deletions were generally shorter in unique piCs compared to common piCs 76 nt vs 148 nt (fig. 3E, Kruskal Wallis p *>*.05). This observation is different from those in *D. melanogaster*, where the length distribution of deletions was similar between unique and common piCs [Srivastav et al., 2023]. However, this supports the idea that unique piCs are emerging *de novo*, considering the larger insertions, which can be associated with recent TE insertions.

### TE enrichment in common and unique piCs

We were interested in determining whether piCs were enriched for particular TE families, and if that enrichment differed between common and unique piCs. We used TE annotations from Repeatmasker to determine the TE composition of piCs in each strain [Smit et al., 2013-2015]. We only considered TE insertions that were at least 500 bp and no more than 20% diverged from the reference sequence (20% is when it is generally considered to be a different TE). We compared TE enrichment between piCs that were common between all five of the genotypes in this study and piCs that were unique to one strain. However, even common piCs can have different TE content in different genotypes, thus we also compared within each strain. We used a binomial test to compare observed TE composition to expected composition based on 1000 random shuffles of TE content.

For both unique and common piCs no TE was depleted compared to the rest of the genome, which is expected given that piCs are enriched for TE sequences (fig. 4). There were many more enriched TEs for common piCs (115-124) compared to unique piCs (9-20) when all genotypes were considered together. In common piCs 102 enriched TEs were shared by at least four genotypes (fig. 4A, B, Supplemental file 4, Figure. 7, 8). In unique piCs six of the enriched TEs were shared by at least four genotypes (*412*, *Doc*, *Jockey-1 DSi*, *Max-Element*, *roo*, and *TRANSIB2*). All of the TEs enriched in unique piCs were non-LTRs or LTRs, and the only DNA transposon close to making this list was *TRANSIB2* which was shared by four genotypes. Among common piCs the enrichment of LTRs and non-LTRs also holds, only 11 and 4 of the TEs shared between genotypes are either a DNA transposon or *Helitron*, respectively. The families which are most commonly enriched in both unique and common piCs were *Gypsy* and *Jockey*. TEs that were enriched in both common and unique piCs of at least four genotypes were the following: *412*, *Doc*, *Jockey-1 Dsim*, *Max* -element, and *TRANSIB2* (Supplemental file 7). Other than *412* these are all different elements than those that most commonly nucleate unique clusters in *D. melanogaster* - (*flea*, *Stalker-2*, *412*, *blood*) [Srivastav et al., 2023]. This most likely is a reflection of the unique invasion history of each species. It is important to note that recent invaders such as *Shellder* would not be expected to be included here as it is a somatic TE which is suppressed by a different pathway centered around the *flamenco* locus. However, it is interesting that neither the *P* -element nor *Spoink* are enriched in piCs, both of which are known recent invaders [Kofler et al., 2015, Scarpa et al., 2025].

### Age of enriched TEs in common and unique piCs

Next, we were interested in investigating the age distribution of these enriched TE insertions. We were interested in this question for the following reasons - for example, if a young TE is enriched in unique piCs, this could suggest that newly invaded TEs are triggering the formation of unique piRNA clusters. A TE with fewer differences from the reference indicates a more recent invasion. We used TEs that were enriched in either unique or common piCs discussed in the previous section to determine the relationship between enrichment and age. For all classes of TEs, unique piCs contained the least diverged insertions (fig. 4C, supplemental file 4, Figure. 9).

Unlike the results of unique piCs, the common piCs and non-piC regions of the genome exhibit more similarity in the mean and median divergence of TE insertions. In fact, non-piC regions have a lower mean and median divergence percentage compared to common piCs for TE insertions belonging to the LTR class in all genotypes (fig. 4C, supplemental file 4, Figure. 9). Across all genotypes LTR class TEs in the non-piC region had an average mean and median divergence of 5.67% and 2.45%, respectively, compared to 6.78% and 4.74% in common piCs. One notable difference between piCs and non-piCs was the range of divergence (difference between the upper and lower quartile), which was much larger in non-piCs for LTR, non-LTR, and DNA TEs than in common piCs. For LTRs the average first and second quartiles ranged from 0.59% to 10.97% in non-piC versus 2.56% to 10.43% in common piCs. For non-LTRs the average first and second quartiles ranged from 1.22% to 9.28% in non-piC versus 3.2% to 6.16% in common piCs. And finally for DNA transposons the average first and second quartiles ranged from 3.27% to 11.11% in non-piCs versus 4.42% to 9.28% in common piCs (fig. 4C, Supplemental file 4, Figure. 9). Using *LNP-01-062* and *SZ45*, we confirmed a significant difference between the average divergence of common piCs compared to unique piCs and non-piC regions for LTRs, DNA transposons, and Helitrons (Wilcoxon rank-sum test, *p*-value *<*0.05). In contrast, the divergence of non-LTR elements was significantly different when common and unique piCs were compared, but not when either was compared with non-piC regions (Wilcoxon rank-sum test, *p*-value *>*0.05).

The same holds true when looking at the TE insertions classified as the top ten TE families by frequency of occurrence. The least diverged TE families belong to unique piCs while the common piCs and non-piC regions of the genome exhibit more similarity in the mean and median divergence of TE insertions (fig. 4D, supplemental file 4 Figure. 10). *Gypsy* (mean 0.72% and median .5%) was the youngest among the top 10 TE families enriched in unique piCs of *SZ129*, this was also true in an additional two genotypes (3 out of 5). Other notably young TE families in this strain included *Pao* (LTR) and *I* (Non-LTR) (fig. 4D). This is largely consistent with previous results which found that LTR retrotransposons tended to be enriched at young piCs.

We also wanted to determine if the difference in TE divergence between common and unique piCs was significant. We selected two genotypes to use as examples, *SZ45* and *LNP-01-062*, and found that TEs were indeed significantly younger in unique piCs (Wilcoxon rank-sum test, p-value *≤* 0.05), suggesting that recent invasions nucleate the formation of new piCs.

## Discussion

In this manuscript, we used population genomic methods to study the evolution of piCs in *D. simulans*, the sister species to *D. melanogaster*. There were two major goals of this study - to determine if the patterns of piC evolution inferred in *D. melanogaster* were consistent in *D. simulans*, and to determine if they were consistent with a ‘trap’ model of TE suppression or a ‘birth and death’ model. This study would not have been possible without the generation of high quality genome assemblies for each *D. simulans* genotype, matched with genotype-specific small RNA libraries. Replication of piRNA libraries was a crucial component of this research, as it verified that the calling of piCs is consistent across collections even after two years of within strain evolution. This also suggests that piRNA clusters are stable over time within a single strain. This is important in the context of this work as it could be argued that given the volume of piCs unique to a single genotype, small piRNA clusters are stochastic byproducts of the piRNA system. For example, any of a number of regions could become piRNA clusters stochastically depending on the organisms development and the pool of maternal piRNA. However, we found that piRNA cluster calls were *<*90% consistent over time, thus even small piRNA clusters are stable within a single strain. This lends weight to our finding that small piRNA clusters tend to be unique to a single strain rather than common across genotypes.

There are aspects of our findings that support the ‘trap’ model of piC evolution, and others that do not. Active TE families are enriched in piCs, which is consistent with the trap model [Bergman et al., 2006, Kofler, 2019]. We also find that larger piCs are enriched for more diverse TE families than small piCs, and they are relatively younger than in non-piC regions. This is consistent with the ‘trap’ model, where piCs represent a library of recent TE activity. We find extensive intra-specific variation in piC activity across genotypes, as has been previously reported [Srivastav et al., 2023, Ellison and Cao, 2020]. It is not clear what causes a piC to lose its piRNA activity, especially given that they are stable within a genotype. It is possible that it is due to structural changes at the piC, but at this point we do not know. However, the frequent formation of piCs throughout the genome is not consistent with the trap model, and it does appear that recent TE insertions nucleate the formation of new piCs [Srivastav et al., 2023]. These newly emerging piCs may increase in frequency and size through drift or selection for TE suppression (though hosts do not appear to be insertion-limited). Over time, these piCs may trap additional TEs and form large clusters such as *42AB*. TEs that no longer produce piRNAs necessary for TE suppression may be lost or become dispensible [Gebert et al., 2021]. It is possible that because piRNA loci are re-established every generation, the piRNA pool may rapidly evolve to lose inactive TE families and thus no longer license the activity of piRNA clusters. Why larger clusters may be lost is unknown, but our work seems more consistent with the ‘birth and death’ model produced by Srivastev (2023) than the ‘trap’ model (for germline piCs) [Bergman et al., 2006, Srivastav et al., 2023]. The largest point of difference is the observation that entirely different families of TEs nucleate young piCs, suggesting it is simply new invasions that trigger piC formation rather than a certain propensity of a TE family.

Differences between *D. melanogaster* and *D. simulans* in the TEs which nucleate unique piRNA clusters likely reflect the invasion history of the species rather than the propensity for any given TE type to nucleate a cluster. In *D. melanogaster* it was predicted that certain types of TEs nucleate clusters, and the most common TEs in small piRNA clusters were *412*, *blood*, *flea*, and *Stalker-2* [Srivastav et al., 2023]. All but one of these TEs is a member of the *mdg3* group of TEs. It was hypothesized that this family of TE might readily license piCs from maternally inherited piRNAs, or they could, as a family, have more of a propensity to produce double stranded RNA and endogenous siRNAs. There has been some recent evidence that endogenous siRNA production precedes piRNA formation, and that maternal inheritance of this piRNA is necessary for the formation of these piCs [Luo et al., 2023].

In *D. simulans* the TEs most associated with unique piCs are *412*, *Doc*, *Jockey-1 Dsim*, *Max* -element, and *TRANSIB2*. One TE is a member of the *mdg3* family (*412*), two are non-LTRs from the *Doc* and *Jockey* families (*Doc*, *Jockey-2 D. simulans*), one belongs to the *BEL-Pao* family (*Max* -element, and one is a DNA element (*TRANSIB2*). Based on what we know about the invasion history of these elements in *D. melanogaster* and *D. simulans*, it seems more likely that the representation of these elements in newer piCs is due to their recent invasion rather than any inherent propensity for producing siRNA. For example, in *D. melanogaster 412* and *blood* invaded in the late 1800s, while the invasion history of *Stalker-2* and *flea* (also known as *Blastopia*) is not evident in historical samples dating back 200 years [Scarpa et al., 2023]. However, other work suggests *Stalker-2* and *Blastopia* are also recent invaders in *D. melanogaster* based on similarity between copies in the genome [Dias and Carareto, 2012]. The invasion history of *D. simulans* is not known beyond modern invasions such as *Shellder* and *Spoink*, however a *Doc* insertion is associated with a recent selective sweep in *D. simulans* [Schlenke and Begun, 2004] and the *Max* -element has been noted as being recently active [Carareto et al., 2014]. *TRANSIB2* in *D. melanogaster* is only present in fragmented copies, and its copy number is much higher in *D. simulans* than in *D. melanogaster* [Gebert et al., 2021, Mohamed et al., 2020]. Furthermore, there were large differences in copy number of *TRANSIB2* between *D. simulans* collected in CA and Africa, which could indicate recent expansion [Signor, 2020]. It is difficult to assess any evidence related to *Jockey-1 D. simulans* due to the ambiguity of its nomenclature. None of these TEs contain an *env* protein, which is consistent with expression in the germline. Regardless, it is most likely that any recent invasion can nucleate a piRNA cluster rather than it being a propensity of a particular type of TE itself.

Originally piCs were described as concentrated regions of the genome that produced large amounts of piRNA and were predicted to be quite conserved, such as *42AB* which produced 20% of total piRNA [Brennecke et al., 2007]. Srivastev (2023) found that even large conserved piCs such as *42AB* can have variable activity in different genotypes of the same species, with some genotypes having almost no production of piRNA from this cluster. Furthermore, Srivastev et al. (2023) found that many piCs are small and unique to single genotypes of *D. melanogaster*. In *D. simulans* we have found the same to be true - the vast majority of piCs are small and unique to a single genotype. However, there has also been considerable evolutionary turnover between *D. melanogaster* and *D. simulans*, in *D. melanogaster* the piCs described as producing the majority of piRNA are not important in *D. simulans*. *42AB* produces less than half a percent of total piRNA in *D. simulans*, and the highest producing clusters are unique to this species. This is an incredible rate of change between two species separated by 2 mya. However, we have shown previously that in *D. melanogaster* 11 TEs have invaded in just the last 200 years [Pianezza et al., 2024, 2025, Scarpa et al., 2025, 2023]. If the *D. melanogaster* genome is co-evolving with TEs as they invade, considerable evolution would have occurred during this time period.

We used much more stringent cutoffs for piCs than in previous work, but piCs still occupied a larger proportion of the genome than in *D. melanogaster*. Different cutoffs will always produce different proportions, but it is quite suggestive that a much larger proportion of the genome is occupied by piCs in *D. simulans* than in *D. melanogaster*. For example, we required that piCs have an RPM of 25, while in work on *D. melanogaster* the requirement was 8 RPKM. It is unclear why a species may have a much larger proportion of piCs than another closely related species. A minimum size for piCs such that transopons can be effectively repressed was predicted to be 3% of the genome. No maximum size for piRNA clusters has been predicted, but it has been theorized that their size could represent a balance between selective forces which increase their size (TE invasions) and those that may decrease their size (likelihood of ectopic recombination, cost of piRNA production) [Kofler, 2020].

In summary, evidence has been accumulating that the ‘trap’ model of piC suppression does not adequately describe the dynamics of germline piC formation in drosophilids. In the somatic support cells of the ovary where *flamenco* is responsible for silencing TEs the ‘trap’ model most likely functions, but in the germline it appears that TEs can nucleate the formation of small piCs. Why a particular insertion may nucleate a piC while another may not is unknown. We do know, based on our work and that of previous studies, that piCs are highly reproducible, suggesting that they are not specified stochastically but rather have some inherent property. Future research on this question will be very informative as to the formation and maintenance of piCs.

## Supplemental Data

Supplemental File 1 contains the restrictive call piC co-ordinates used in the study. Supplemental File 2 provides the discontinued proTRAC call results. Supplemental File 3 provides data of overlap annalysis between proTRAC and restrictive calls. Supplemental File 4 contains all supplementary figures referenced throughout the manuscript. Supplemental File 5 contains the replicate restrictive call piC co-ordinates used to assess reproducibility and genotype specific piRNA cluster stability. Supplemental File 6 presents piRNA count distribution among piCs across all genotypes. Supplemental file 7 provides the TE enrichment annalysis information of common and unique piCs among all genotypes included in the study.

## Acknowledgments

S.S. would like to acknowledge S. and F. Emery for help in preparing the manuscript, as well as A. Palmier for inspiration.

## Author contributions

P.N. performed all of the data analysis, interpretation, and figure presentation. S.S. concieved of the study. P.N. and S.S. prepared the manuscript.

## Funding

This work was supported by the National Science Foundation Established Program to Stimulate Competitive Research grants NSF-EPSCoR-1826834 and NSF-EPSCoR-2032756 to SS and NIH R35GM155272 to SS.

## Conflicts of Interest

The author(s) declare(s) that there is no conflict of interest regarding the publication of this article.

## Data Availability

All data used in this manuscript are publicly available under NCBI Genome BioProject PRJNA907284 for the the genome assemblies and NCBI SRA PRJNA913883 for the small RNA sequencing.

## Notes

### Competing Interest Statement

The authors have declared no competing interest.

