## Supplementary material for "Rapid evolution and comparative analysis of piRNA clusters in *D. simulans*": Suplemental File 4

### Supplementary Figures

November 2025

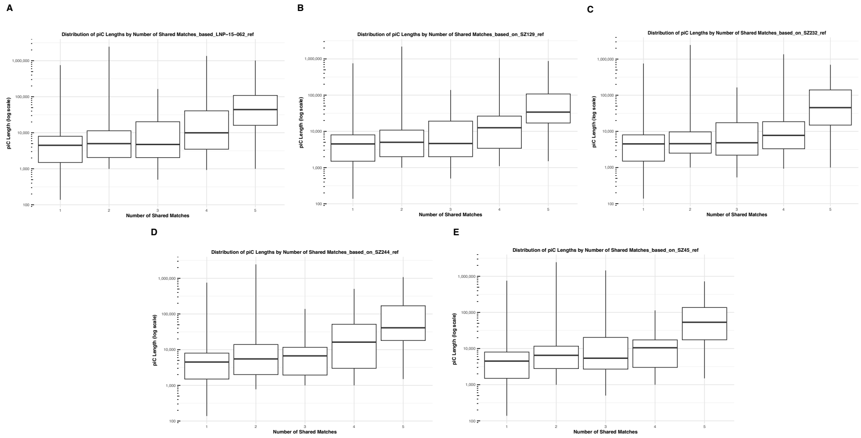


Figure 1: Pi-rna cluster length distribution by number of genotypes shared by a piC, where panels A-E represent results when each genotype was designated as a reference genome.

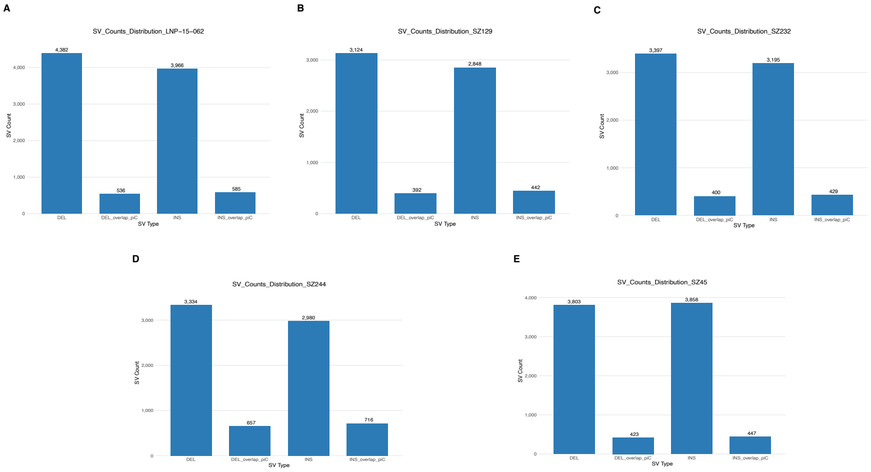


Figure 2: SV counts distribution of insertions and deletions between piC overlap and non-overlap, where panels A-E represent results when each genotype was designated as a reference genome

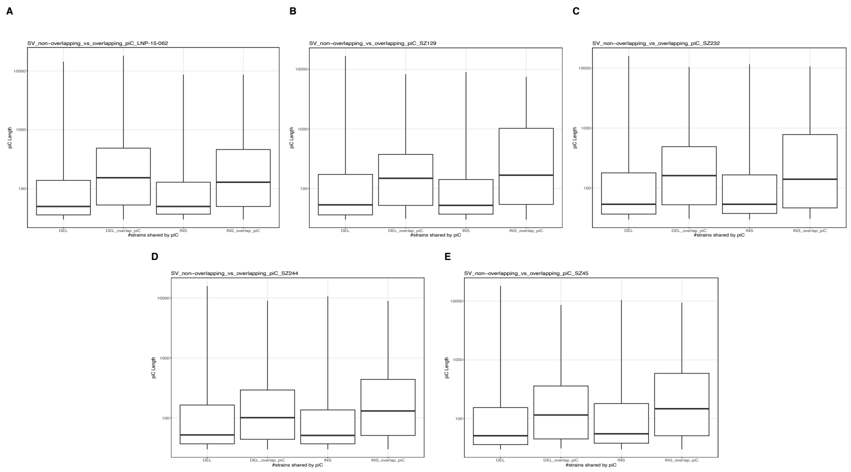


Figure 3: SV length distribution of insertions and deletions overlapping and non-overlapping with piCs, where panels A-E represent results when each genotype was designated as a reference genome.

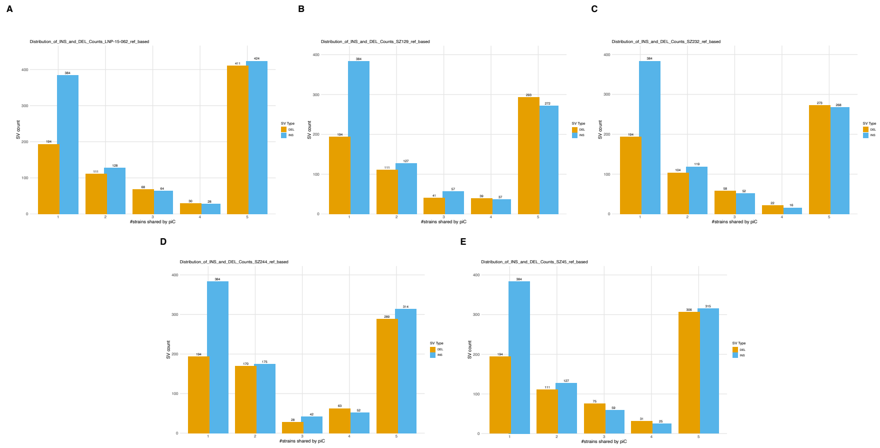


Figure 4: Count of insertions and deletions overlapping piCs grouped by the number of genotypes they are shared by, where panels A-E represent results when each genotype was designated as a reference genome.

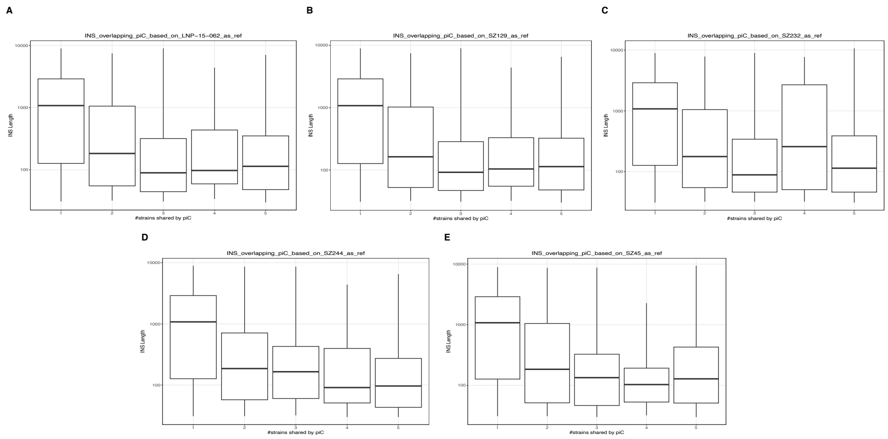


Figure 5: length distribution of insertions overlapping with piCs grouped by the number of genotypes they are shared by, where panels A-E represent results when each genotype was designated as a reference genome.

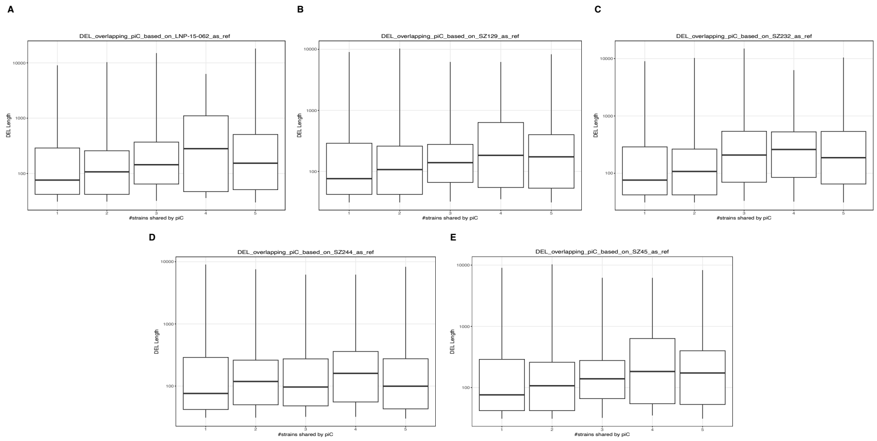


Figure 6: Length distribution of deletions overlapping with piCs grouped by the number of genotypes they are shared by, where panels A-E represent results when each genotype was designated as a reference genome.

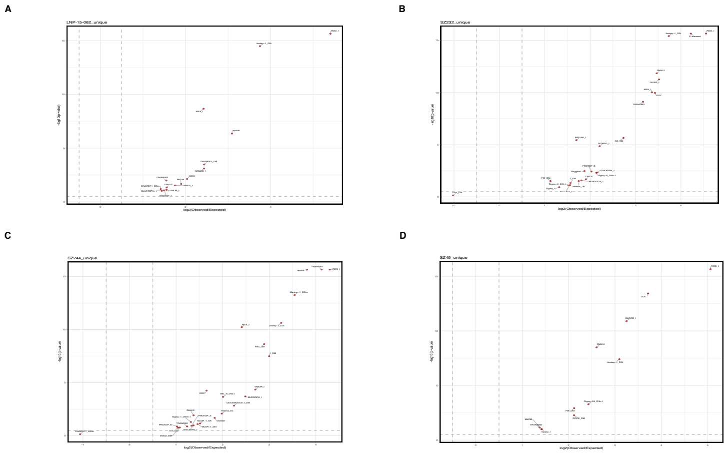


Figure 7: Enrichment analyses of TEs in unique piCs using random shuffling. Results shown for genotypes (A) *LNP-15-062* , (B) *SZ232* , (C) *SZ244* , (D) *SZ45*.

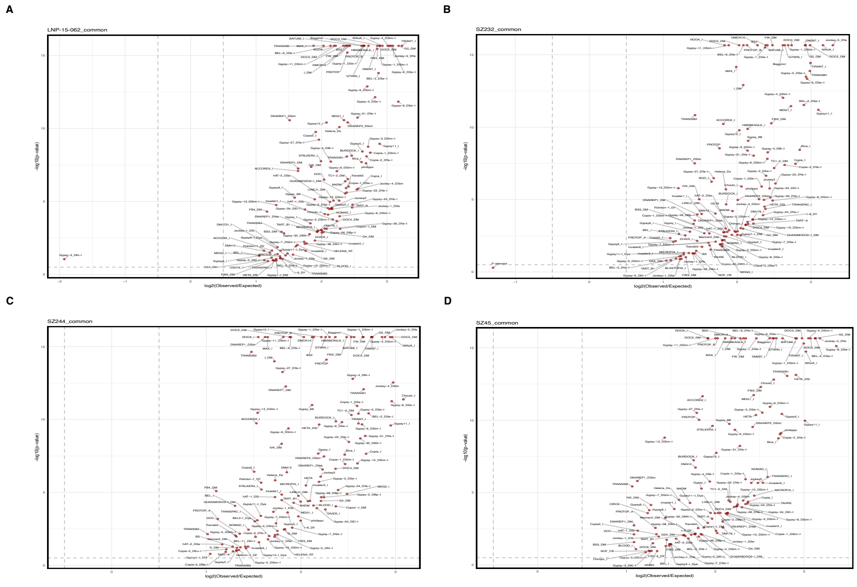


Figure 8: Enrichment analyses of TEs in common piCs using random shuffling. Results shown for genotypes (A) *LNP-15-062* , (B) *SZ232* , (C) *SZ244* , (D) *SZ45*.

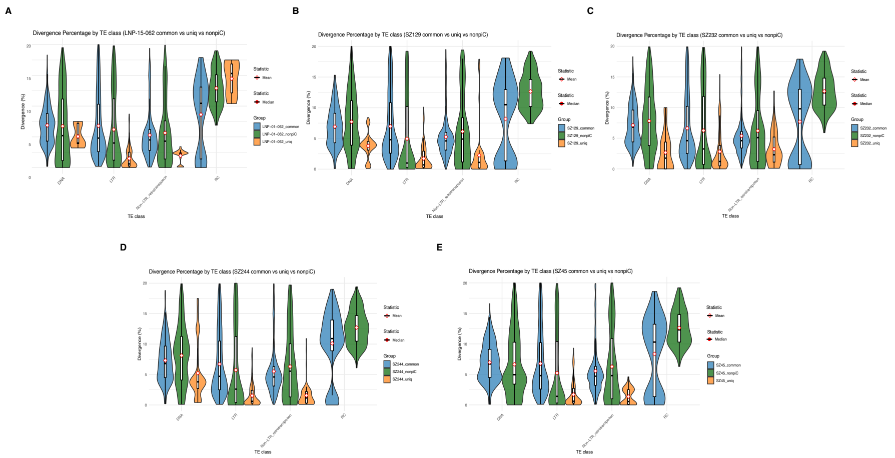


Figure 9: Divergence percentage distribution of TE classes in common piC, non-piC, and unique piCs. Results shown for genotypes (A) *LNP-15-062* , (B) *SZ129* , (C) *SZ232* , (D) *SZ244* , (E) *SZ45*.

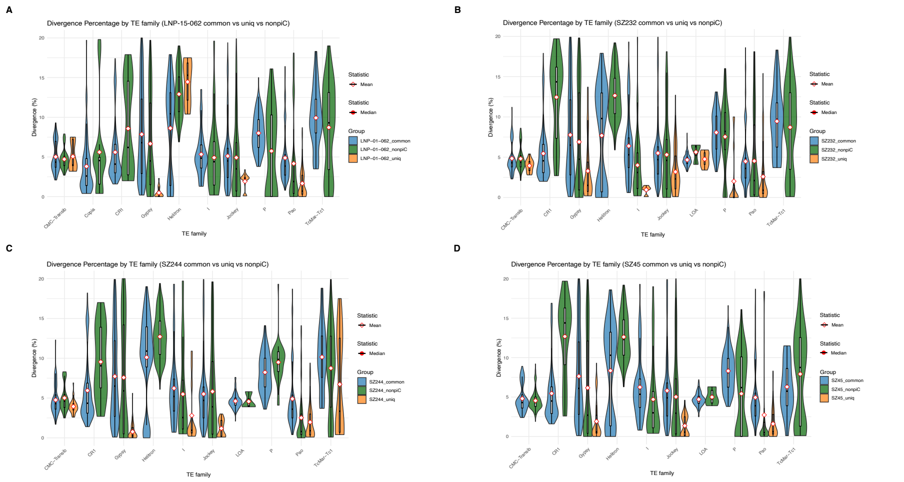


Figure 10: Divergence percentage distribution of top 10 TE families in common piC, non-piC , and unique piCs. Results shown for genotypes (A) *LNP-15-062* , (B) *SZ232* , (C) *SZ244*, (D) *SZ45*.
